# Individual and temporal variation in pathogen load predicts long-term impacts of an emerging infectious disease

**DOI:** 10.1101/392324

**Authors:** Konstans Wells, Rodrigo K. Hamede, Menna E. Jones, Paul A. Hohenlohe, Andrew Storfer, Hamish I. McCallum

**Affiliations:** Environmental Futures Research Institute, Griffith University, Brisbane QLD 4111, Australia; School of Biological Sciences, University of Tasmania, Private Bag 55, Hobart, Tasmania 7001, Australia; Institute for Bioinformatics and Evolutionary Studies, Department of Biological Sciences, University of Idaho, Moscow, ID 83844, USA; School of Biological Sciences, Washington State University, Pullman, WA 99164-4236 USA

**Author notes:** E-mail addresses (following order of authorship).

**Keywords:** disease burden, long-periodicity oscillation, population viability, Tasmanian devil, transmissible cancer, wildlife health

## Abstract

Emerging infectious diseases increasingly threaten wildlife populations. Most studies focus on managing short-term epidemic properties, such as controlling early outbreaks. Predicting long-term endemic characteristics with limited retrospective data is more challenging. We used individual-based modelling informed by individual variation in pathogen load and transmissibility to predict long-term impacts of a lethal, transmissible cancer on Tasmanian devil (*Sarcophilus harrisii*) populations. For this, we employed Approximate Bayesian Computation to identify model scenarios that best matched known epidemiological and demographic system properties derived from ten years of data after disease emergence, enabling us to forecast future system dynamics. We show that the dramatic devil population declines observed thus far are likely attributable to transient dynamics. Only 21% of matching scenarios led to devil extinction within 100 years following devil facial tumour disease (DFTD) introduction, whereas DFTD faded out in 57% of simulations. In the remaining 22% of simulations, disease and host coexisted for at least 100 years, usually with long-period oscillations. Our findings show that pathogen extirpation or host-pathogen coexistence are much more likely than the DFTD-induced devil extinction, with crucial management ramifications. Accounting for individual-level disease progression and the long-term outcome of devil-DFTD interactions at the population-level, our findings suggest that immediate management interventions are unlikely to be necessary to ensure the persistence of Tasmanian devil populations. This is because strong population declines of devils after disease emergence do not necessarily translate into long-term population declines at equilibria. Our modelling approach is widely applicable to other host-pathogen systems to predict disease impact beyond transient dynamics.

## Introduction

Emerging infectious diseases most often attract attention because their initial impacts on host populations are frequently severe (de Castro and Bolker 2005, Smith et al. 2009). Following the initial epidemic and transient dynamic behaviour, long-term outcomes include pathogen fadeout, host extinction, or long-term endemicity with varying impacts on the host population size (Hastings 2004, Benton et al. 2006, Cazelles and Hales 2006). Predicting which of these long-term outcomes may occur on the basis of initial transient dynamics is very challenging and conclusions about possible disease effects on population viability based on early epidemic dynamics can be misleading.

Nevertheless, predicting the long-term consequences of an infectious disease as early as possible in the emergence process is important for management. If the disease has a high likelihood of ultimately leading to host extinction, then strategies such as stamping out infection by removing all potentially infectious individuals may be justifiable, despite short-term impacts on the host species and ethical considerations (McCallum and Hocking 2005). Resource-intensive strategies such as establishing captive breeding populations protected from disease or translocating individuals to locations separated from infected populations may also be justified (McCallum and Jones 2006). In contrast, if impacts are transitory, then a preferred strategy may be to avoid interference to allow a new long-term endemic disease state or pathogen extinction to be reached as quickly as possible (Gandon et al. 2013). Longer-term evolutionary processes can operate to ultimately reduce the impact of the disease on the host population (Fenner 1983, Kerr 2012), and inappropriate disease management strategies may slow down evolution of both host and pathogen.

Models of infectious diseases in the early stages of emergence typically focus on estimating *R_0_*, the number of secondarily infected individuals when one infected individual is introduced into a wholly susceptible population (Lloyd-Smith et al. 2005). This is a key parameter for devising strategies to limit invasion or control an outbreak because it allows the estimation of vaccination or removal rates necessary to eradicate disease. However, by definition, it does not include density dependent factors and is therefore sometimes insufficient to predict the long-term consequences of disease introduction into a new population.

Most existing models for infectious disease are based around compartmental Susceptible – Exposed – Infected – Recovered epidemiological models (S-E-I-R), which rely on a strict assumption of homogeneity of individuals within compartments (Anderson and May 1991). There is a parallel literature for macroparasitic infections, which assumes both a stationary distribution of parasites amongst hosts and that parasite burden is determined by the number of infective stages the host has encountered (Anderson and May 1978). For many pathogens, pathogen load on (or inside) an individual typically changes following infection as a result of within-host processes, causing temporal shifts in transmission and host mortality rates. For example, the volume of transmissible tumours on Tasmanian devils (*Sarcophilus harrisii*) increases through time, with measurable impacts on survival (Wells et al. 2017) and likely temporal increases in transmission probability to uninfected devils that bite into the growing tumour mass (Hamede et al. 2013). Similarly, increasing burden of the amphibian chytrid fungus *Batrachochytrium dendrobatidis* on individual frogs after infection limits host survival, with important consequences for disease spread and population dynamics (Briggs et al. 2010, Wilber et al. 2016). Burdens of the causative agent of white nose syndrome, *Pseudogymnoascus destructans*, which threatens numerous bat species in North America, similarly increase on most individuals during the period of hibernation (Langwig et al. 2015). The additional time dependence introduced by within-host pathogen growth can have a major influence on the dynamics of host-pathogen interactions as uncovered by nested models that link within- and between-host processes of disease dynamics (Gilchrist and Coombs 2006, Mideo et al. 2008). Such dynamics are poorly captured by conventional compartmental and macroparasite model structures. Thus, connecting across the scales of within- and between-host dynamics remains a key challenge in understanding infectious disease epidemiology (Gog et al. 2015).

Here we develop an individual-based model to explore the long-term impact of devil facial tumour disease (DFTD), a transmissible cancer, on Tasmanian devil populations. DFTD is a recently emerged infectious disease, first detected in 1996 in north-eastern Tasmania (Hawkins et al. 2006). It is caused by a clonal cancerous cell line, which is transmitted by direct transfer of live tumour cells when devils bite each other (Pearse and Swift 2006, Jones et al. 2008, Hamede et al. 2013). DFTD is nearly always fatal and largely affects individuals that are otherwise the fittest in the population (Wells et al. 2017). Population declines to very low numbers concomitant with the frequency-dependent transmission of DFTD led to predictions of devil extinctions, based on compartmental epidemiological models (McCallum et al. 2009, Hamede et al. 2012).

Fortunately, the local devil extinctions predicted from these early models have not occurred (McCallum et al. 2009). There is increasing evidence that rapid evolutionary changes have taken place in infected devil populations, particularly in loci associated with disease resistance and immune response (Epstein et al. 2016, Pye et al. 2016, Wright et al. 2017). Moreover, we recently reported that the force of infection (the rate at which susceptible individuals become infected) increases over a time period of as long as six years (~3 generations) after initial local disease emergence and that the time until death after initial infection may be as long as two years (Wells et al. 2017). Therefore, despite high lethality, the rate of epidemic increase appears to be relatively slow, prompting predictive modelling of population level impacts over time spans well beyond those covered by field observations.

In general, there are three potential long-term outcomes of host-pathogen interactions: host extinction, pathogen extirpation, and host-pathogen coexistence. To determine the likelihood of each of these outcomes in a local population of Tasmanian devils, we used individual-based simulation modelling (**Figure 1**) and pattern matching, based on ten years of existing field data, to project population trajectories for Tasmanian devil populations over 100 years following DFTD introduction.

**Figure 1.**
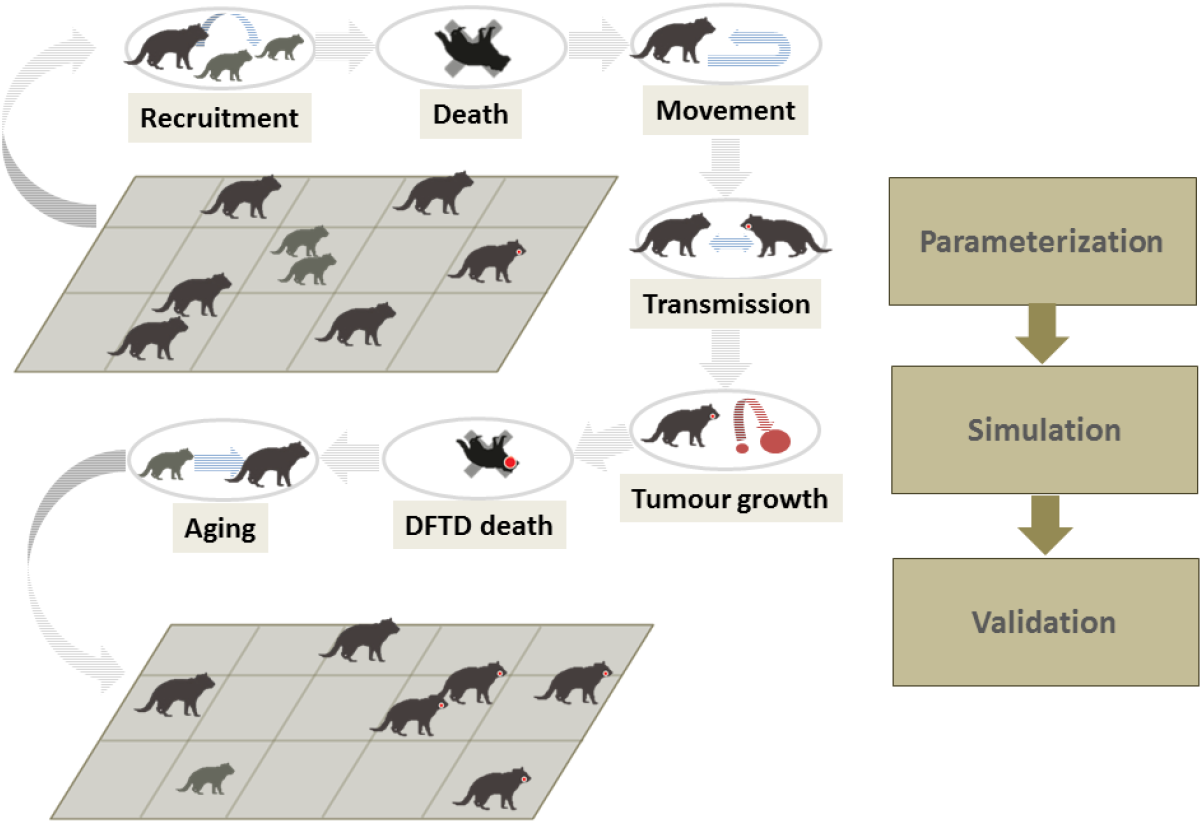
Illustrative overview of the individual-based model to explore long-term population changes of a Tasmanian devil population burdened with devil facial tumour disease (DFTD). Individuals are distributed in a study area. For every weekly time step seven different processes are modelled, namely 1) the possible recruitment of young from females (conditional on young survival during previous weaning time), 2) possible death independent of disease status, 3) movement of individuals away from their home range centre, 4) behavioural interaction between nearby individuals that may result in the transmission of DFTD, 5) growth of DFTD tumours, 6) death of individuals resulting from DFTD, 7) aging of individuals.

## Materials and methods

### Model framework

We implemented a stochastic individual-based simulation model of coupled Tasmanian devil (*Sarcophilus harrisii*) demography and devil facial tumour disease (DFDT) epidemiology. A full model description with overview of design, concept, and details (Grimm et al. 2006) can be found in SI Appendix 1. In brief, we aimed to simulate the impact of DFTD on Tasmanian devil populations and validate 10^6 model scenarios of different random input parameters (26 model parameters assumed to be unknown and difficult or impossible to estimate from empirical studies, see Supporting Information **Table S1**) by matching known system level properties (disease prevalence and population structure) derived from a wild population studied over ten years after the emergence of DFTD (Hamede et al. 2015). In particular, running model scenarios for 100 years prior to, and after the introduction of DFTD, we explored the extent to which DFTD causes devil populations to decline or become extinct. Moreover, we aimed to explore whether input parameters such as the latency period of DFTD or the frequencies of disease transmission between individuals of different ages can be identified by matching simulation scenarios to field patterns of devil demography and disease prevalence.

Entities in the model are individuals that move in weekly time steps (movement distance *θ*) within their home ranges and may potentially engage in disease-transmitting biting behaviour with other individuals (**Fig 1**). Birth-death processes and DFTD epidemiology are modelled as probabilities according to specified input parameter values for each scenario. In each time step, processes are scheduled in the following order: 1) reproduction of mature individuals (if the week matches the reproductive season), 2) recruitment of juveniles into the population, 3) natural death (independent of DFTD), 4) physical interaction and potential disease transmission, 5) growth of tumours, 6) DFTD-induced death, 7) movement of individuals, 8) aging of individuals.

The force of infection *λ_i,t_*, i.e. the probability that a susceptible individual *i* acquires DFTD at time *t* is given as the sum of the probabilities of DFTD being transmitted from any interacting infected individual *k* (with *k* ∈ 1…*K*, with *K* being the number of all individuals in the population excluding *i*):

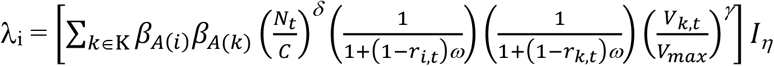

Here, the disease transmission coefficient is composed of the two factors *β_A(i)_* and *β_A(k)_*, each of which accounts for the age-specific interaction and disease transmission rate for individuals *i* and *k* according to their age classes *A*. *N_t_* is the population size at time *t;* the scaling factor *δ* accounts for possible increase in interactions frequency with increasing population size if *δ* > 0. The parameter *r_i,t_* is a Boolean indicator of whether an individual recently reproduced and *ω* is a scaling factor that determines the difference in *λ_i,t_* resulting from interactions of reproductively active and non-reproducing individuals. *V_k,t_* is the tumour load of individual *k*, *V_max_* is the maximum tumour load, and *γ* is a scaling factor of how *λ_i,t_* changes with tumour load of infected individuals. The parameter *I_η_* is a Boolean indicator of whether two individuals are located in a spatial distance < *η* that allows interaction and disease transmission (i.e. only individuals in distances < *η* can infect each other). We considered individuals as ‘reproductively active’ (*r_i,t_*=1) for eight weeks after a reproduction event.

DFTD-induced mortality *Ω_size_* accounts for tumour size, while tumour growth was modelled as a logistic function with the growth parameter α sampled as an input parameter. We allowed for latency periods *τ* between infection and the onset of tumour growth, which was also sampled as an input parameter. We assumed no recovery from DFTD, which appears be very rare in the field (Pye et al. 2016).

Notably, sampled scaling factor values of zero for *δ, ω*, and *γ* correspond to model scenarios with homogeneous interaction frequencies and disease transmission rates independent of population size, reproductive status and tumour load, respectively, while values of *η* = 21 km assume that individuals can infect each other independent of spatial proximity (i.e. individuals across the entire study area can infect each other). The sampled parameter space included scenarios that omitted *i*) effects of tumour load on infection and survival propensity, *ii*) effect of spatial proximity on the force of infection between pairs of individuals and *iii*) both effects of tumour load and spatial proximity, in each of 1,000 scenarios. This sampling design was used to explicitly assess the importance of modelling individual tumour load and space use for accurately representing the system dynamics.

### Model validation and summary

To resolve the most realistic model structures and assumptions from a wide range of possibilities and to compare simulation output with summary statistics from our case study (a devil population at West Pencil Pine in western Tasmania) (Wells et al. 2017), we used likelihood-free Approximate Bayesian Computation (ABC) for approximating the most likely input parameter values, based on the distances between observed and simulated summary statistics (Toni et al. 2009). We used the ‘neuralnet’ regression method in the R package *abc* (Csillery et al. 2012). Prediction error was minimized by determining the most accurate tolerance rate and corresponding number of scenarios considered as posterior through a subsampling cross validation procedure as implemented in the *abc* package. For this, leave-one-out cross validation was used to evaluate the out-of-sample accuracy of parameter estimates (using a subset of 100 randomly selected simulated scenarios), with a prediction error estimated for each input parameter (Csillery et al. 2012); this step facilitates selecting the most accurate number of scenarios as a posterior sample. However, we are aware that none of the scenarios selected as posterior samples entirely represents the true system dynamics. We identified *n* = 122 scenarios (tolerance rate of 0.009, Supporting Information **Figure S1**) as a reasonable posterior selection with minimized prediction error but sufficiently large sample size to express uncertainty in estimates. The distribution of summary statistics was tested against the summary statistics from our case study as a goodness of fit test, using the ‘gfit’ function in the *abc* package (with a p-value of 0.37 indicating reasonable fit, see Supporting Information **Figure S2, S3**).

We generated key summary statistics from the case study, in which DFTD was expected to have been introduced shortly before the onset of the study (Hamede et al. 2015), and a pre-selection of simulation scenarios, in which juveniles never comprised > 50% of the population, DFTD prevalence at end of 10-year-period was between 10 and 70%, and the age of individuals with growing tumours was ≥ 52 weeks. Hereafter, we refer to ‘prevalence’ as the proportion of free-ranging devil individuals (animals ≥ 35 weeks old) with tumours of sizes ≥ 0.1 cm^3^; we do so to derive a measure of prevalence from simulations that is comparable to those inferred from field data. Summary statistics were: 1) mean DFTD prevalence over the course of 10 years, 2) mean DFTD prevalence in the 10^th^ year only, 3) autocorrelation value for prevalence values lagged over one time step, 4) three coefficient estimates of a cubic regression model of the smoothed ordered difference in DFTD prevalence (fitting 3^rd^ order orthogonal polynomials of time for smoothed prevalence values using the loess function in R with degree of smoothing set to α = 0.75), 5) phase in seasonal population fluctuations, calculated from sinusoidal model fitted to the number of trappable individuals in different time steps, 6) regression coefficient of a linear model of the changing proportions of individuals ≥ 3years old in the trappable population over the course of ten years. Summary statistics for the simulations were based on the 37 selected weekly time steps after the introduction of DFTD that matched the time sequences of capture sessions in the case study, which included records in ca. three months intervals (using the first 30 time steps only for population sizes, as the empirical estimates from the last year of field data may be subject to data censoring bias). Overall, these summary statistics aimed to describe general patterns rather than reproducing the exact course of population and disease prevalence changes over time, given that real systems would not repeat themselves for any given dynamics (Wood 2010). Additionally, unknown factors not considered in the model may contribute to the observed temporal changes in devil abundance and disease prevalence.

As results from our simulations, we considered the posterior distributions of the selected input parameters (as adjusted parameter values according to the ABC approach utilised) and calculated the frequency and timing of population or disease extirpation from the 100 years of simulation after DFTD introduction of the selected scenarios. All simulations and statistics were performed in R version 3.4.3 (R Development Core Team 2017). We used wavelet analysis based on Morlet power spectra as implemented in the R package *WaveletComp* (Roesch and Schmidbauer 2014) to identify possible periodicity at different frequencies in the time series of population sizes (based on all free-ranging individuals) for scenarios in which DFTD persisted at least 100 years.

## Results

For scenarios that best matched empirical mark-recapture data, 21% of scenarios (26 out of 122) led to devil population extirpation in timespans of 13 – 42 years (mean = 21, SD = 8; ~7–21 generations) after introduction of DFTD (**Figure 2**). In contrast, the disease was lost in 57% of these scenarios (69 out of 122), with disease extirpation taking place 11 – 100 years (mean = 29, SD = 22) post-introduction (**Figure 2**). Loss of DFTD from local populations therefore appears to be much more likely than devil population extirpation, given no other factor than DFTD reducing devil vital rates. Moreover, fluctuations in host and pathogen after the introduction of DFTD exhibited long-period oscillations in most cases (**Figure 3**). In the 27 selected scenarios in which DFTD persisted in populations for 100 years after disease introduction, population size 80–100 years after disease introduction was smaller and more variable (mean = 137, SD = 36) than population sizes prior to the introduction of DFTD (mean = 285, SD = 3; **Figure 4**). The average DFTD prevalence 80–100 years after disease introduction remained < 40% (mean = 14%, SD = 4%; **Figure 4**). Most wavelet power spectra of these scenarios showed long-period oscillations over time periods between 261 – 1040 weeks (corresponding to 5 – 20 years) (electronic supplementary material, Supporting Information **Figure S4**).

**Figure 2.**
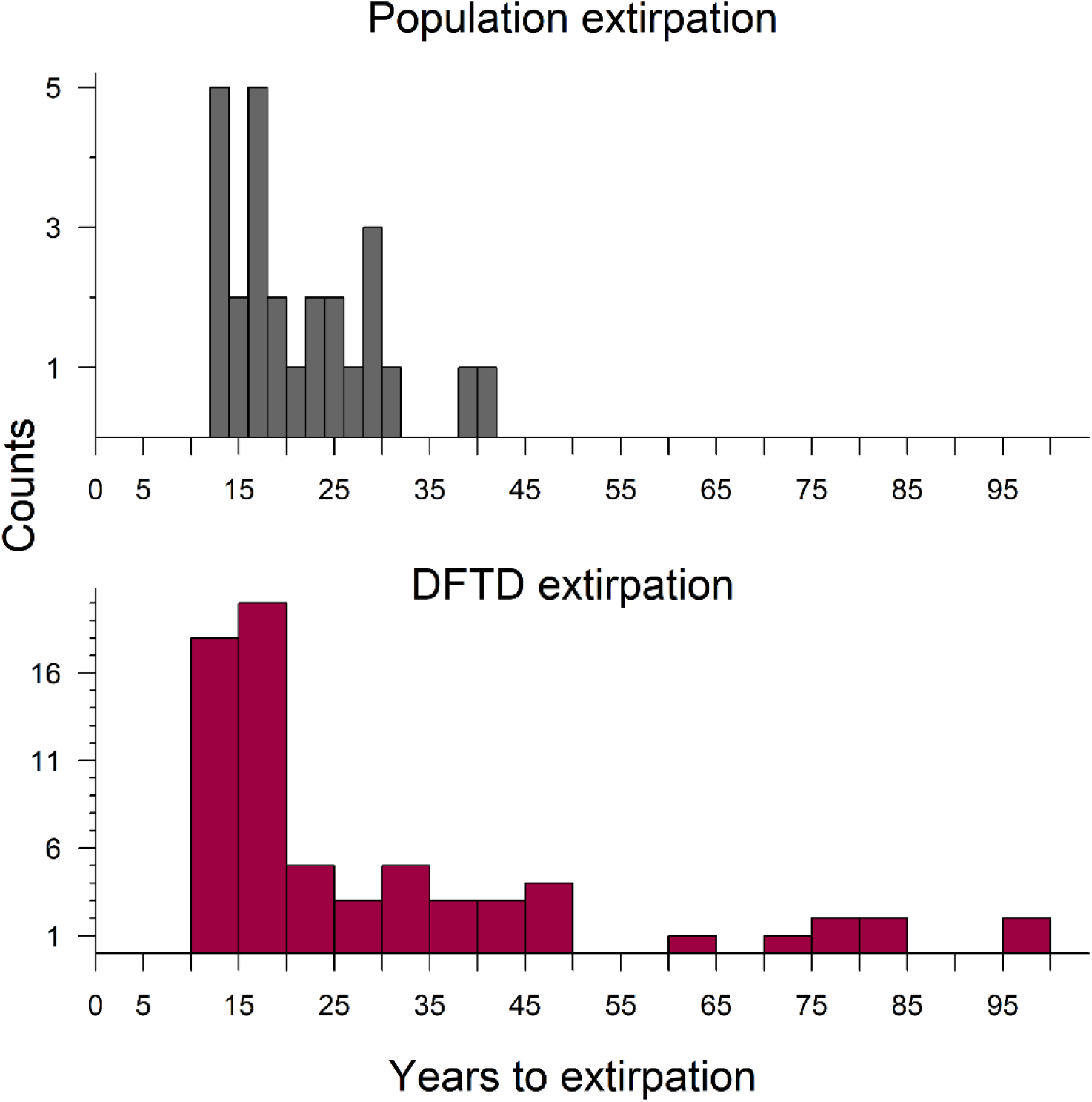
Frequency distributions of timespans of devil extirpation (upper panel) and devil facial tumour disease (DFTD) extirpation presented as years after the introduction of the disease into populations. Number of plotted scenarios correspond to those for which extirpation events were recorded (26 and 69 out of 122 posterior samples, respectively).

**Figure 3.**
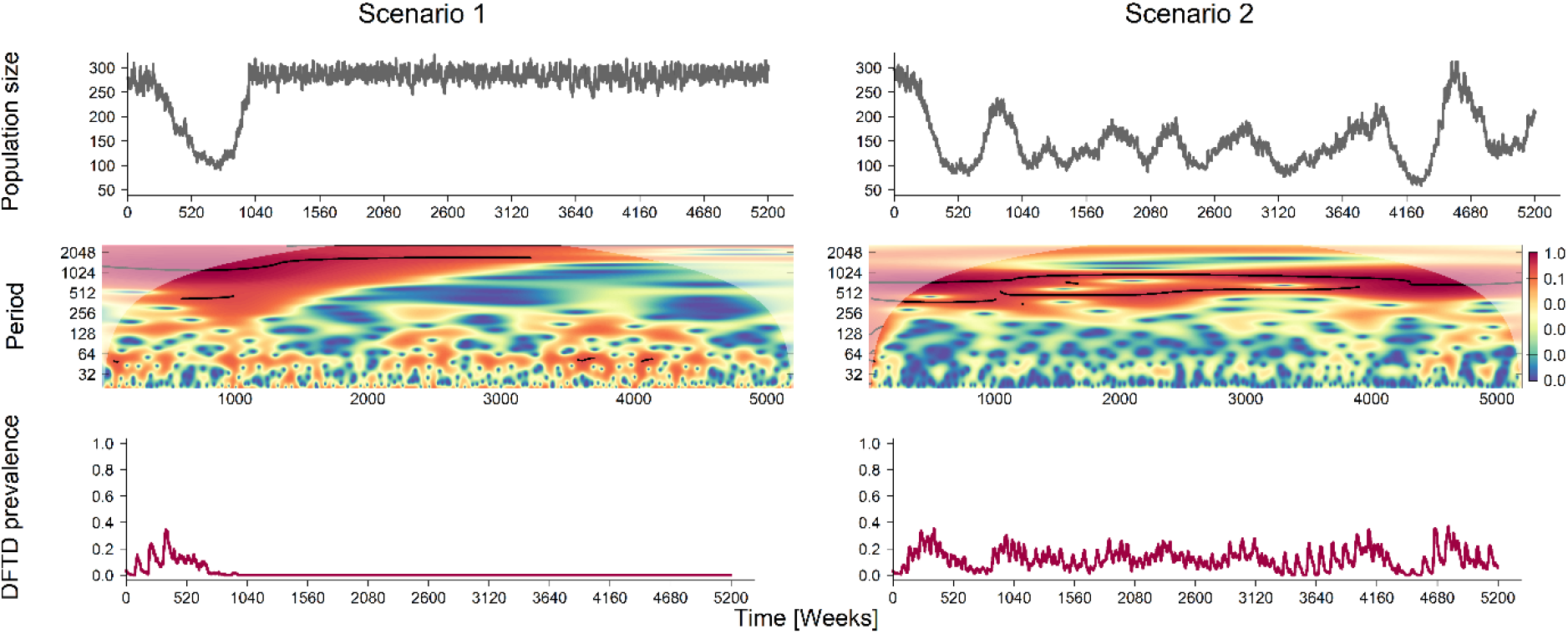
Examples of long-term devil and tumour dynamics. Scenario 1 is an example of DFTD extirpation, and Scenario 2 is an example of coexistence. The upper panels show the summarized population sizes (free-ranging individuals ≥ 35 weeks old) over 100 years (5,200 weeks) of simulations after the introduction of DFTD in the population, middle panels show the respective wavelet power spectra, based on Morlet wavelet analysis. Red colours in the power spectra show periodicity (measured in weeks) of highest intensity with ridges (black lines) at frequencies often > 500 weeks. Lower panels show the prevalence of DFTD (growing tumour ≥ 0.1 cm^3^) in the respective population.

**Figure 4.**
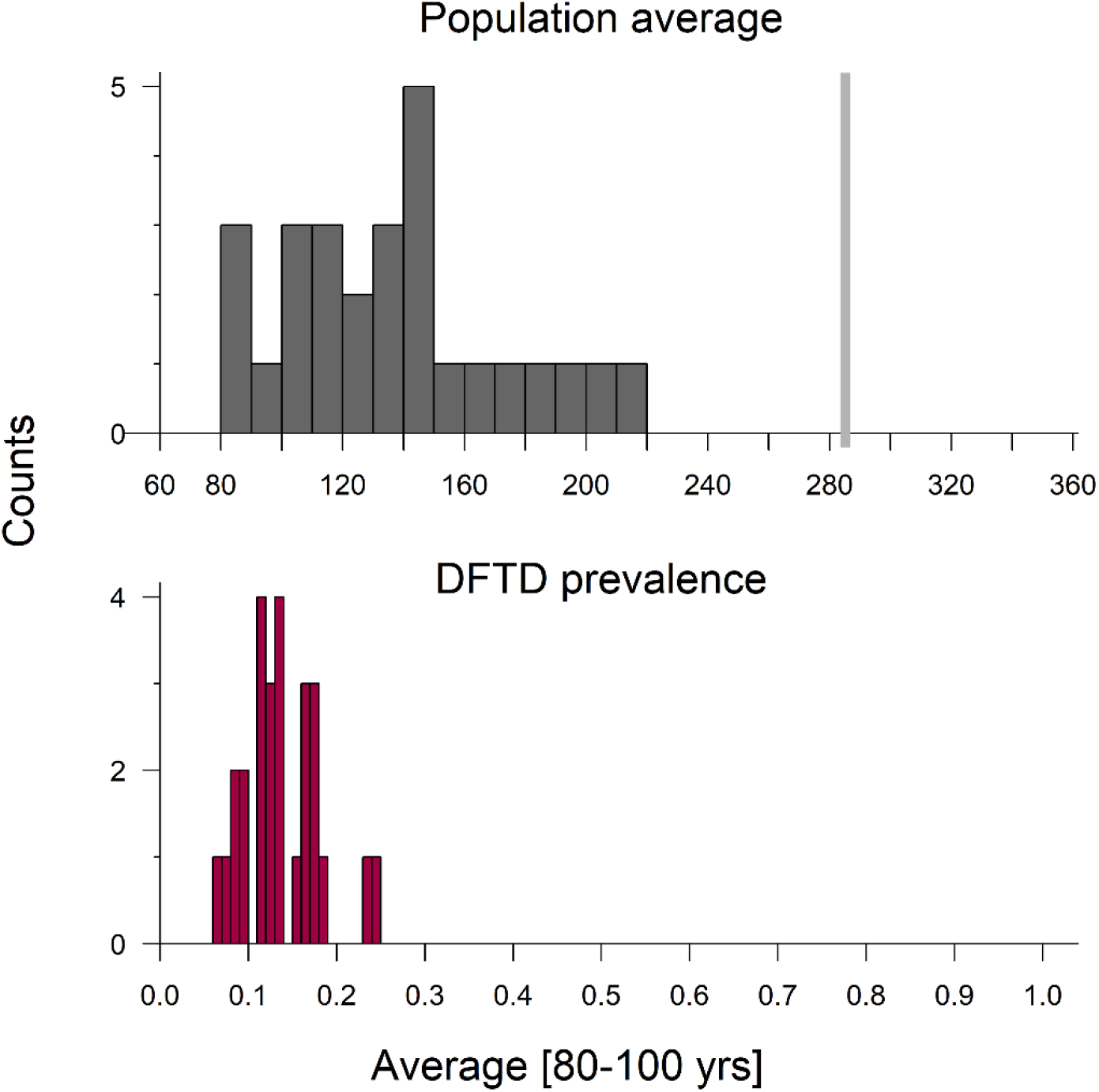
Frequency distributions (count) of mean devil populations sizes (x axis, upper panel) and devil facial tumour disease (DFTD) prevalence (x axis, lower panel) 80–100 years after disease introduction for those scenarios (n = 27) in which DFDT persisted for at least 100 years. The light-grey vertical line in the upper panel indicates the mean population sizes of simulated populations over 100 years prior to disease introduction.

Inference of input parameters was only possible for some parameters, whereas 95% credible intervals for most of the posterior distributions were not distinguishable from the (uniformly) sampled priors. Notably, the posterior mode for the latency period (*τ*) was estimated as 50.5 weeks (95% credible interval 48.5 – 52.6 weeks, for unadjusted parameters values the 95% was 22.9 – 94.3 weeks), providing a first estimate of this latent parameter from field data (Supporting Information **Figure S5, Table S2**). The posterior of the DFTD-induced mortality factor (odds relative to un-diseased devils) for tumours < 50 cm^3^ (*Ω_<50_*) was constrained to relatively large values (electronic supplementary material, Supporting Information **Figure S5**), supporting empirical estimates that small tumours are unlikely to cause significant mortality of devils. Posterior distributions of weekly movement distances (*θ*) and the spatial distance over which disease-transmitting interactions took place (*η*), in turn, allowed no clear estimates of these parameters (electronic supplementary material, Supporting Information **Figure S5**). Notably, the 122 scenarios selected as posteriors all explicitly accounted for the effect of tumour load on infection and survival, while 90% of selected scenarios included spatial proximity of individuals as influencing disease transmission (i.e. selected scenarios comprised 110 models that included both the effect of tumour load and spatial proximity, while 12 models included tumour load but not spatial proximity).

## Discussion

Our results suggest that DFTD will not necessarily cause local Tasmanian devil extinction or even long-term major declines, whereas the extirpation of DFTD or coexistence/endemicity is much more likely. In cases where DFTD persists in local devil populations in the long-term, oscillations with relatively long periods (5–20 years, corresponding to 2–10 generations) appear likely. These predictions are starkly different from those derived from previous compartmental models, which considered all devils with detectable tumours to be equally infectious and assumed exponentially distributed time delays. These models predicted extinction (McCallum et al. 2009), as did models with more realistic gamma distributed time delays or with delay-differential equations that incorporated field-derived parameter estimates of transmission and mortality rates (Beeton and McCallum 2011). These previous models, however, differ also from our approach in that they ignore spatial structure and do not account for the uncertainty in unknown parameters such as disease-induced mortality and disease transmission rates.

The predictions from our individually-based model, derived from 10 years of observational data at our case study site (West Pencil Pine), are consistent with observations now emerging from long-term field studies of the dynamics of Tasmanian devils and DFTD (Lazenby et al. 2018). No Tasmanian devil population has yet become extinct – and populations persist, albeit in low numbers, where disease has been present the longest (e.g., at wukalina/Mount William National Park and at Freycinet, where DFTD emerged, respectively, at least 21 and 17 years ago) (Epstein et al. 2016). Also, a considerable decline in DFTD prevalence has been observed in recent years at Freycinet (Sebastien Comte, unpublished data). These study sites did not contribute to the fitting of our model and at least to some extent constitute an independent validation and test of the model predictions. Our modelling results suggest that observed population dynamics of devils and DFTD do not require evolutionary changes, although there is evidence of rapid evolution in disease-burdened devil populations (Epstein et al. 2016) similar to rapid evolution in other vertebrates when subjected to intense selection pressure (Christie et al. 2016, Campbell-Staton et al. 2017).

One of the differences between earlier models and those we present here is the inclusion of tumour growth, with mortality and transmission rates that depend on individual disease burden. Inclusion of burden-dependent dynamics results in additional and qualitatively different time delays than those incorporated in previous models. Tumours take time to grow before they have a major impact on host survival and become highly infectious (Hamede et al. 2017, Wells et al. 2017). This slows the spread of DFTD and its impact on devil population fluctuations. It also means that parameters estimated from field data, without taking tumour growth into account, do not adequately represent the system dynamics (McCallum et al. 2009). Our models suggest that documented dramatic population declines during the first 10 years or so of the DFTD epizootic may represent just the first peak of a classical epidemic (Bailey 1975). Long-term predictions from our models suggest, however, that DFTD is a slow burning disease with population changes governed by long-term oscillations.

It is well known, both from simple Lotka-Volterra models and from a range of empirical studies, that consumer–resource interactions have a propensity to cycle, driven by the time delays inherent in these systems (Murdoch et al. 2003). Disease burden-dependent demographic and epidemiological parameters, together with burden growth within the host, add additional time delays, both lengthening any oscillations and increasing the likelihood that they will be maintained in the longer term. Apparently, such time-delays increase the probability of host-pathogen coexistence, similar to predator-prey dynamics, rather than host or pathogen extirpation. Grounded in theory and a reasonable body of modelling studies of other wildlife diseases, disease-induced population extinction appears to be more generally an exception rather than the rule, unless host populations are very small, or unless there are reservoir species that are tolerant of infection (de Castro and Bolker 2005).

The approach we apply here – coupling the flexibility of individual-based models to account for heterogeneity in disease burden and space use with Approximate Bayesian Computation to match model outcomes with available empirical evidence - offers considerable potential for making predictions regarding the population dynamics for other emerging diseases (Toni et al. 2009, Beaumont 2010, Johnson and Briggs 2011, Wells et al. 2015). A fundamental problem in applying modelling approaches to forecast the outcome of emerging infectious disease epidemics is the need to estimate parameter values based on empirical data derived from the relatively early stages of an epizootic, in the absence of retrospective knowledge (Heesterbeek et al. 2015, Ferguson et al. 2016). Examples include estimating *R_0_* for SARS (Lipsitch et al. 2003) and for the 2014–2015 Ebola epidemic in West Africa (Whitty et al. 2014, WHO Ebola Response Team 2014) among others (LaDeau et al. 2011). In most of these cases, the objective is to estimate parameters associated with the growth phase of the epidemic to assess the effectiveness of interventions such as vaccination. The task we have addressed in this paper is even more challenging – seeking to predict the long-term endemic behaviour of a pathogen that is currently still in the early stages of emergence. We suggest that management efforts to maintain devil populations in the face of DFTD should be guided by our changing understanding of the long-term dynamics of the DFTD epidemic.

Management efforts in wild populations that solely aim to combat the impact of DFTD can be counterproductive if they disrupt long-term eco-evolutionary dynamics that may eventually lead to endemicity with stable devil populations. Our ability to predict future outcomes in the absence of management actions require some caution as we cannot fully exclude the possibility that DFTD can cause local population extinctions once populations are small, warranting future research. While our findings emphasize the importance of accounting for individual tumour load for accurate prediction and epidemiological modelling of DFTD dynamics, our inability to uncover the exact role of devil spatial proximity on disease transmission means that further research is necessary to understand relevant factors in disease spread.

The key management implication of our model is that “heroic” management interventions are unlikely to be necessary to ensure persistence of Tasmanian devil populations. Given more information on immune-related or genetic variation in resistance, the model could be modified to assess the value of interventions such as vaccination or reintroduction of captive reared animals. At the same time, we believe that any management actions should be subject to rigorous quantitative analysis to explore possible long-term impacts. In particular, allocating resources and scientific endeavours to the management of wildlife diseases such as DFTD should not disguise the fact that sufficiently large and undisturbed natural environments are a vital prerequisite for wildlife to persist and eventually cope with perturbations such as infectious diseases without human intervention.

## Acknowledgments

The study was funded by NSF grant DEB 1316549 and NIH grant R01 GM126563–01 as part of the joint NIH-NSF-USDA Ecology and Evolution of Infectious Diseases Program, the Australian Research Council Discovery (DP110102656), Future Fellowship (FT100100250), DECRA (DE17010116) and Linkage (LP0561120, LP0989613) Schemes, the Fulbright Scholarship Scheme, the Ian Potter Foundation, the Australian Academy of Science (Margaret Middleton Fund), two grants from the Estate of W.V. Scott, the National Geographic Society, the Mohammed bin Zayed Conservation Fund, the Holsworth Wildlife Trust and three Eric Guiler Tasmanian Devil Research Grants through the Save the Tasmanian Devil Appeal of the University of Tasmania Foundation. We are grateful for all support during field work underpinning this modelling study: M. and C. Walsh, Discovery Holidays Parks-Cradle Mountain National Park, and Forico Pty Ltd provided logistic support and many volunteers helped with data collection (published previously). Comments from Samuel Alizon and anonymous reviewers improved previous drafts.

## Authors’ contribution

K.W. conceived the idea of this study, carried out the analysis and wrote the first draft. All authors interpreted results and contributed to revisions. All authors gave final approval for publication.

